# Activation of a Latent Epitope Causing Differential Binding of Anti-Neutrophil Cytoplasmic Antibodies to Proteinase 3

**DOI:** 10.1101/549063

**Authors:** Marta Casal Moura, Gwen E. Thompson, Darlene A. Nelson, Lynn A. Fussner, Amber M. Hummel, Dieter E. Jenne, Daniel Emerling, Wayne Volkmuth, Fernando C. Fervenza, Cees G.M. Kallenberg, Carol A. Langford, W. Joseph McCune, Peter A. Merkel, Paul A. Monach, Philip Seo, Robert F. Spiera, E. William St. Clair, Steven R. Ytterberg, John H. Stone, William H. Robinson, Yuan-Ping Pang, Ulrich Specks, the WGET and RAVE-ITN Research Groups

## Abstract

**Objective:** Proteinase 3 (PR3) is the major antigen for anti-neutrophil cytoplasmic antibodies (ANCAs) in the systemic autoimmune vasculitis, granulomatosis with polyangiitis (GPA). PR3 anti-neutrophil cytoplasmic antibodies (PR3-ANCAs) recognize different epitopes on PR3. We aimed to study the effect of mutations on PR3 antigenicity.

**Methods:** The recombinant PR3 variants, iPR3 which is clinically used to detect PR3-ANCAs and iHm5 which contains three point mutations in Epitope 1 and 5 generated for epitope mapping studies, immunoassays and serum samples from patients enrolled in ANCA-associated vasculitis (AAV) clinical trials were used to screen the differential PR3-ANCA binding. Selective binding was determined by inhibition experiments.

**Results:** Rather than a reduced binding of PR3-ANCAs to iHm5, we found substantially increased binding of the majority of PR3-ANCAs to iHm5 compared with iPR3. A monoclonal ANCA (moANCA518) from a patient with GPA was found to selectively bind to iHm5 within the mutation-free Epitope 3 and distant from the point mutations of iHm5 contained in Epitope 1 and 5. Binding of iPR3 to monoclonal antibody MCPR3-2 also induced recognition by moANCA518.

**Conclusion:** The preferential binding of PR3-ANCAs from patients like the selective binding of moANCA518 to iHm5 is conferred by increased antigenicity of Epitope 3 on iHm5. This can also be induced on iPR3 when it is captured by monoclonal antibody MCPR-2. This previously unrecognized characteristic of PR3-ANCA interactions with its target antigen has implications for studying antibody-mediated autoimmune diseases, understanding of variable performance characteristics of immunoassays and design of potential novel treatment approaches.

## 1. Introduction

Anti-neutrophil cytoplasmic autoantibody (ANCA)-associated vasculitis (AAV) comprises a group of systemic small-vessel vasculitis syndromes including granulomatosis with polyangiitis (GPA), microscopic polyangiitis (MPA), and eosinophilic granulomatosis with polyangiitis (EGPA) (1). The two major target antigens for the ANCAs in vasculitis are proteinase 3 (PR3) and myeloperoxidase (MPO) (2-4). Testing for the presence of ANCAs has become indispensable in the evaluation of patients suspected of having AAV (5). Patients with PR3-targeting ANCAs (PR3-ANCAs) are at higher risk for relapses than patients with myeloperoxidase-targeting ANCAs (MPO-ANCAs). In addition, PR3-ANCA positivity and rising titers following treatment portend relapses, particularly for patients with renal disease and other disease manifestations of capillaritis (6-8). The binding of PR3-ANCAs to PR3 has many well-documented pro-inflammatory effects, and PR3-ANCAs are thought to play a pathogenic role for the development of necrotizing vasculitis (9-11).

The oligoclonal PR3-ANCAs from patients with GPA are known to bind to different epitopes of the folded PR3 antigen, whereas denatured PR3 or improperly folded recombinant PR3 generated in non-mammalian expression systems do not bind PR3-ANCAs reliably (12-14). Consistently, studies of continuous epitope mapping of PR3-ANCAs using oligopeptides have generated inconclusive results (15-18).

In this context, we performed instead discontinuous epitope mapping of anti-PR3 monoclonal antibodies (moAbs) and PR3-ANCAs using human-murine chimeric recombinant PR3 molecules with surface epitope-specific point mutations to obtain mechanistic insights into how interactions of PR3 with its environment during inflammation are modified by PR3-ANCAs and potentially targetable by therapeutics (19). Herein, the iPR3 mutant represents the most prevalent wild-type PR3 conformation (Val103) and contains the Ser195Ala mutation at the active site (**Fig. 1**) to avoid potential enzymatic degradation of ANCAs or PR3-capturing monoclonal antibodies (moAbs) by PR3 in immunoassays or cytotoxicity when recombinant PR3 is expressed in the human kidney epithelial cells 293 (HEK293) (20-23). Because iPR3 has the same folded conformation of the wild-type mature PR3, it has been used for two decades as a standard PR3 antigen for sensitive and specific PR3-ANCA detection by immunoassay (7, 22, 24-29).

**Fig. 1.**
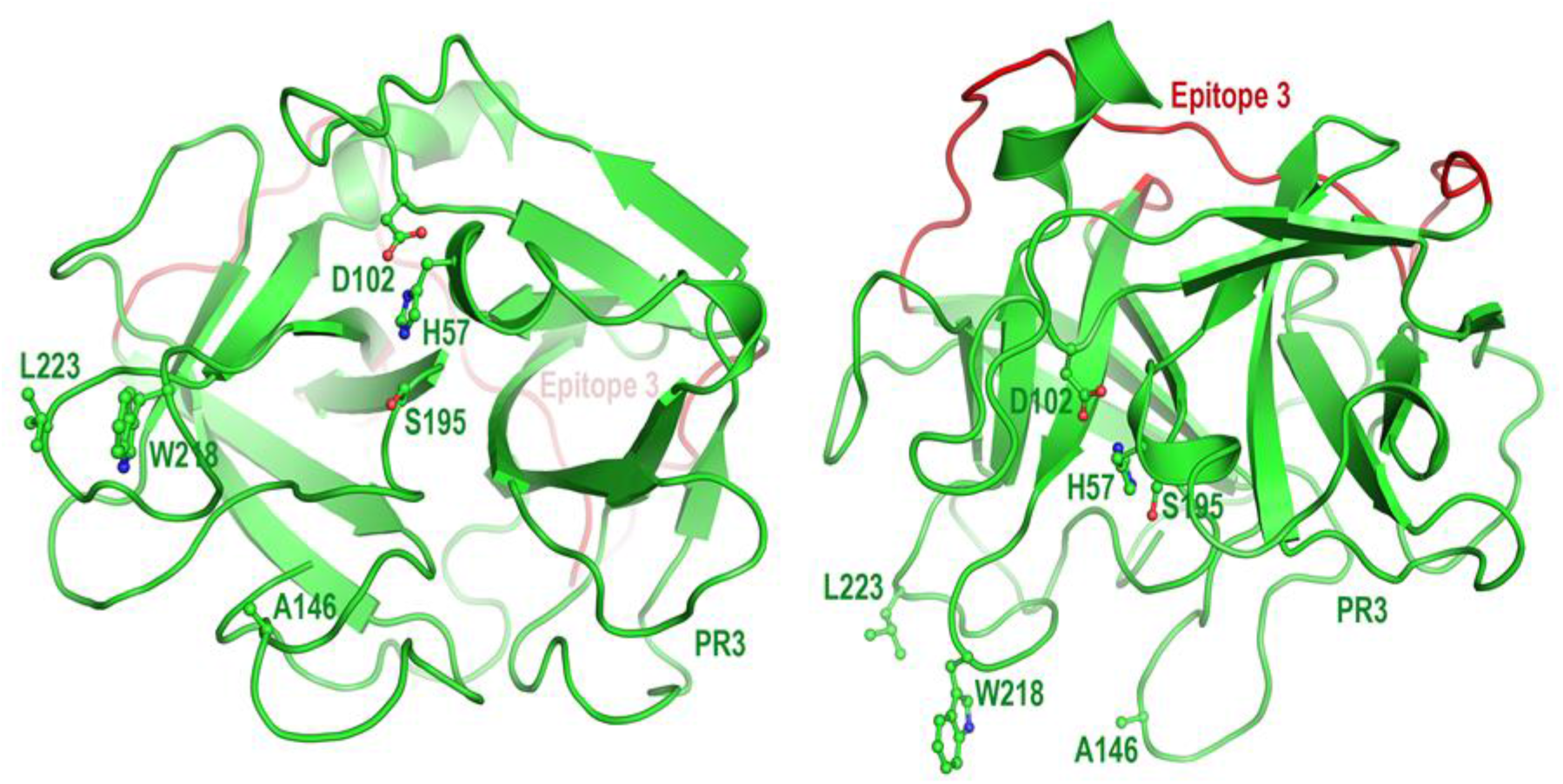
Cartoon model of human proteinase 3 (PR3)*. The left panel shows PR3 in the standard orientation facing the active site pocket; the right panel shows PR3 with an approximately 90 degree rotation. The catalytic triad of this neutrophil serine protease comprises His57, Asp102, and Ser195. Ala146, Trp218, and Leu223 in PR3 shown here are replaced by their hydrophilic murine counterparts (Thr146, Arg218, and Gln223) in iHm5. Shown in red is Epitope 3 on the opposite side of the PR3 structure from the side where the mutations were introduced. (*Generated using PyMOL v. 1.7.0.3, Schroedinger, LLC; amino acid numbering is based on Fujinaga M. et al. J. Mol. Biol. 1996; 261:267-78)

In our epitope mapping study, we developed a human-murine chimeric mutant of iPR3 (iHm5; formerly referred to as Hm5 (19)) to investigate the involvement of Epitope 5 on PR3— an epitope defined by binding of a group of moAbs including MCPR3-7—in the binding of PR3 to neutrophil membranes (19, 30, 31). Because iHm5 has its three hydrophobic/aromatic residues (Ala146, Trp218, and Leu223; **Fig. 1**) of human PR3 replaced by their murine hydrophilic counterparts (Thr146, Arg218, and Gln223), we expected that the PR3-ANCAs would show reduced binding to the mutated epitopes on iHm5 relative to iPR3. Instead, we serendipitously identified a human moAB (moANCA518) derived from a patient with GPA that recognized iHm5, but not iPR3, and demonstrated that this preferential recognition of iHm5 by moANCA518 was caused by the increased mobility of Epitope 3 leading to the remote activation of a latent epitope on PR3 by the distant mutation (32).

Informed by this background, we conducted the present study using large well-defined patient cohorts to determine (i) the scope of this preferential binding of PR3-ANCAs to iHm5, (ii) the potential utility of iHm5 as an *in vitro* antigen for PR3-ANCA detection, (iii) the epitope(s) involved in this enhanced antigenicity of iHm5 for PR3-ANCAs, and (iv) whether remote activation of a latent epitope could also be induced by binding of moAbs to distant epitopes of iPR3.

## 2. Material and methods

Reagents were obtained from Sigma (St. Louis, MO) unless specified otherwise. The HEK293 cell line used for the expression of recombinant PR3 mutants was obtained from ATCC (Rockville, MD).

### 2.1. Recombinant PR3 mutants

The cDNA constructs coding for iPR3 and iHm5 and their expression in HEK293 cells were described in detail elsewhere (19, 26). Both mutants carry a carboxy-terminal c-myc-peptide extension and a poly-His peptide extension for anchoring in solid phase immunoassays and purification using nickel columns from GE Healthcare (Chicago, IL) as previously described and specified below (19, 26-28, 33).

### 2.2. Immunoassays

For *Western blots*, PR3 mutants were loaded (1 μg/lane) onto 12% Tris-HCl gels from BioRad (Hercules, CA). Protein samples were not reduced, and SDS gel electrophoresis was performed at 180 volts for 35 minutes. Proteins were transferred from gels to nitrocellulose membranes. Membranes were subsequently washed with TBS buffer, blocked for 45 minutes at room temperature (RT) with TBS / 0.2% non-fat dry milk. Next, membranes were washed twice with TBS / 0.1% Tween 20. Monoclonal antibodies, diluted to 0.5–1.0 μg/mL as indicated, were incubated on the membrane overnight at 4 °C. Next, membranes were washed twice with TBS / 0.1% Tween 20 and incubated with goat anti-human or anti-mouse IgG HRP conjugates, diluted to 1:20,000, for 20 minutes at RT. Membranes were washed again and developed with the Pierce ECL Western Blotting Substrate kit from Thermo Fisher Scientific (Waltham, MA) according to package insert, and exposed as indicated.

For the *direct ELISA* experiments, Maxisorp® from Invitrogen (Carlsbad, CA) plates were coated with iHm5 or iPR3, 1.0 μg/mL, in NaHCO_3_ (pH 9.5), overnight at 4 ºC. In between steps, plates were washed three times with 200 μL of PBS with 0.05% Tween 20 (10 mM sodium phosphate, 0.15 M NaCl, 0.05% Tween-20, and pH 7.5). The plates were then blocked with 200 μL of PBS with 1% Tween 20 and 10% BSA diluted to 1:10 for 2 hours, at RT and protected from direct light exposure. moANCA518 or epitope specific anti-PR3 moAbs (1.0 μg/mL) were diluted to 1:20 in PBS with 1% Tween 20 and 10% BSA and used as primary antibodies. The antibody binding to the PR3 mutants was probed with HRP-conjugated anti-human IgG antibody (1:250 dilution) incubated for 1 hour at RT. As substrate, 100 μL of 3,3’,5,5’-tetramethylbenzidine (TMB, Thermo Fisher Scientific®) was developed for 15 minutes and stopped with 100 μL 2 N H_2_SO_4_. The absorbance in optical density (OD) was measured using spectrophotometry at 450 nm. Results are expressed as the net absorbance in OD after subtraction of the absorbance readings of the background wells coated with NaHCO_3_ only. For the inhibition experiments, epitope specific anti-PR3 moAbs (see below) were coated to Maxisorp® plates at concentrations of 2.0–4.0 μg/mL in NaHCO_3_ overnight at 4 ºC to capture the PR3 mutants.

The *anchor ELISA* method for IgG PR3-ANCAs was described previously for IgA PR3-ANCAs (33) and is conceptually similar to the capture ELISA utilizing the *C*-terminal cmyc-peptide extension of a PR3 mutant (26, 27). Either the purified PR3 mutants or culture media supernatants from PR3 mutant expressing 293 cell clones diluted in IRMA buffer (50 mM Tris, 0.1 M NaCl, pH 7.4, and 0.1% BSA) were incubated in Pierce® nickel-coated plates from Thermo Fisher Scientific (Waltham, CA) for 1 hour at RT; the background wells were incubated with serum-free media instead (26, 27, 33). Plates were washed three times with TBS wash buffer (20 mM Tris, 500 mM NaCl, pH 7.5, and 0.05% Tween 20) in between steps. Serum samples were diluted 1:20 in TBS buffer containing 0.5% BSA and incubated with the antigen for 1 hour at RT. PR3-ANCAs were detected after incubation for 1 hour at RT with alkaline phosphatase-conjugated goat anti-human IgG (1:10,000 dilution). When required, IgA and IgM PR3-ANCAs were detected using alkaline phosphatase-conjugated goat anti-human IgA, (1:2,000 dilution) and goat anti-human IgM (μ-chain specific)-alkaline phosphatase antibody (1:20,000), respectively. *P*-Nitrophenyl phosphate was used as substrate at a concentration of 1 mg/mL. The net absorbance was obtained by spectrophotometry at 405 nm after 30 minutes of exposure. The assay’s cut-off value for a positive result was defined as the mean net absorbance obtained from 30 normal control serum samples plus 2 standard deviations of the mean. The net absorbance cut-off values for IgG PR3-ANCA detection using iPR3 and iHm5 were determined as 0.105 and 0.107 OD, respectively.

The *MCPR3-2 capture ELISA* method for PR3-ANCAs detection was described previously and used here to assess the binding of the moANCA518 to iHm5 and iPR3(24). In brief, microtiter wells (Immulon 1B, Thermo, Milford, MA) were incubated with 100 µL of 4 mg/mL MCPR3-2, in NaHCO_3_ (pH 9.5), at 4ºC overnight. After washing three times with TBS wash buffer (20 mM Tris, 500 mM NaCl, pH 7.5, and 0.05% Tween-20), 100 µL of culture media supernatants from PR3 mutant expressing 293 cell clones diluted in IRMA buffer (0.05 mM Tris, 0.1 M NaCl, pH 7.4, 0.1% BSA) were incubated in the wells for 1h at RT. Plates were washed three times with TBS wash buffer (20 mM Tris, 500 mM NaCl, pH 7.5, and 0.05% Tween-20) in between steps. Control wells were incubated in parallel with buffer alone. The moANCA518 solution (1 μg/mL) was prepared in TBS (20 mM Tris, 500 mM NaCl, pH 7.5) with 0.5% BSA and used as the primary antibody in incubation with the antigen for 1h at RT. The moANCA518 binding to the antigen was detected after incubation for 1 hour at RT of alkaline phosphatase conjugated goat anti-human IgG (Sigma, A-9544) antibody in a 1:10,000 dilution in TBS (20 mM Tris, 500 mM NaCl, pH 7.5) with 0.5% BSA. P-nitrophenyl phosphate was used as substrate at a concentration of 1 mg/mL. The net OD was obtained by spectrophotometry at 405 nm after 30 minutes of exposure.

### 2.3. Serum and plasma samples

The 30 serum samples from normal donors used for the determination of the normal (negative) range of the anchor ELISA for PR3-ANCA detection using the PR3 mutants as antigen were obtained from the Clinical Immunology Laboratory of Mayo Clinic, Rochester, MN. Three hundred serum samples from healthy octogenarians used for the specificity analyses were obtained from the Mayo Clinic Biospecimen repository. No patient identifiers or clinical data about these individuals were available to the investigators. The use of these serum samples for this study was approved by the Mayo Clinic Institutional Review Board.

Serum samples from patients with ANCA-associated vasculitis used in this study were collected during the Wegener’s Granulomatosis Etanercept Trial (WGET) and during the Rituximab versus Cyclophosphamide for ANCA-Associated Vasculitis trial (RAVE). Details of the WGET and RAVE protocols, patient characteristics and trial results were described elsewhere (34-36). Participants of both trials had provided written consent for the use of their serum samples in ancillary studies.

For inhibition experiments with antigen-binding fragments (Fab) we used PR3-ANCA positive plasma samples from patients with GPA collected for a biospecimen repository approved by the Mayo Clinic Institutional Review Board. We had previously shown that the agreement of PR3-ANCA levels determined by immunoassay in matched serum and plasma samples is excellent (37).

### 2.4. Generation of moANCA518

DNA barcode-enabled sequencing of the antibody repertoire was performed on plasmablasts derived from 5 participants in the RAVE study at baseline, at time of remission and at time of subsequent relapse as described for rheumatoid arthritis and Sjögren syndrome, and this is described elsewhere (32, 38, 39). The generated recombinant human monoclonal antibodies were tested for reactivity with ANCA target antigens, including MPO (33), human neutrophil elastase (HNE) (40-42), iPR3, and iHm5 in parallel, by the anchor ELISA using recombinant antigens carrying a *C*-terminal poly-His tag.

### 2.5. Monoclonal antibodies and antigen-binding fragments

The moAb PR3G-2 (targeting Epitope 1) was a gift from Prof. C.G.M. Kallenberg from the University of Groningen, and the moAb WGM2 (targeting Epitope 3) was purchased from Hycult Biotech Inc. (Wayne, PA) (19, 43, 44). The moAbs MCPR3-2 (targeting Epitope 4), MPR3-3 (targeting Epitope 3) and MCPR3-7 (targeting Epitope 5) that were generated by Dr. Specks group at Mayo Clinic were described and characterized in detail elsewhere (19, 24, 26).

The Pierce™ Fab Preparation Kit from Thermo Fisher Scientific was used to generate the Fabs from 1.0 mg of a moAb (moANCA518, WGM2, MCPR3-2, MCPR3-3, or MCPR3-7) following the manufacturer’s protocol.

### 2.6. Statistical analysis

IBM^®^ SPSS^®^ Statistics for MacOS, version 25 (IBM, Armonk, NY, USA) was used to calculate the means and standard errors of 3–5 repeat experiments and to compare the means between groups with the two-tailed paired *t*-test.

## 3. Results

### 3.1. Preferential binding of PR3-ANCAs to iHm5 over iPR3

Using the net absorbance as a measure of the reactivity of a serum sample with an antigen, we found that the reactivities of IgG PR3-ANCA positive serum samples with iHm5 were either equal or higher than those with iPR3 in an anchor ELISA (**Fig. 2**). Of the 178 serum samples obtained from the WGET participants at enrollment, 148 had previously tested positive for PR3-ANCA in at least one of several immunoassays for PR3-ANCAs (34). By the anchor ELISA, 144 samples tested positive for IgG PR3-ANCAs with iHm5, and 135 were positive with iPR3 (**Fig. 3 & Table 1**). Of the 135 samples that reacted with both antigens, 108 (80%) had higher reactivities with iHm5 than with iPR3, and 41 (30%) of the samples had reactivities with iHm5 that were more than double the reactivities with iPR3 (**Fig. 3 & Table 1**).

**Table 1.** PR3-ANCA Reactivities with iHm5-Val^103^ and iPR3-Val^103^.

| The WGET study* |  |  |
| --- | --- | --- |
| 148 PR3-ANCA-positive samples* |  |  |
| iHm5-Val<sup>103</sup> positive<br>n = 144 | iPR3-Val<sup>103</sup> positive<br>n = 135 |  |
| Comparison of reactivity in 135 samples positive for both antigens |  |  |
| iHm5-Val<sup>103</sup> > iPR3-Val<sup>103</sup><br>n = 108<br>80% | iHm5-Val<sup>103</sup> = iPR3-Val<sup>103</sup><br>n = 26<br>19% | iHm5-Val<sup>103</sup> < iPR3-Val<sup>103</sup><br>n = 1<br>1% |
| The RAVE trial |  |  |
| 129 PR3-ANCA-positive samples** |  |  |
| iHm5-Val<sup>103</sup> positive<br>n = 128 | iPR3-Val<sup>103</sup> positive<br>n = 126 |  |
| Comparison of reactivity in 126 samples positive for both antigens |  |  |
| iHm5-Val<sup>103</sup> > iPR3-Val<sup>103</sup><br>n = 89<br>71% | iHm5-Val<sup>103</sup> = iPR3-Val<sup>103</sup><br>n = 34<br>27% | iHm5-Val<sup>103</sup> < iPR3-Val<sup>103</sup><br>n = 3<br>2% |

Comparison of the net absorbance values of the PR3-ANCA reactivity with iHm5-Val<sup>103</sup> and iPR3-Val<sup>103</sup>. In 71-80% of the samples, a higher PR3-ANCA recognition of iHm5-Val<sup>103</sup> was documented. Additionally, in 19-27% of the patients, iHm5-Val<sup>103</sup> and iPR3-Val<sup>103</sup> were equally recognized. Very few patients displayed higher recognition of iPR3-Val<sup>103</sup>: 1 in the RAVE trial and 3 in the WGET. [When the confidence interval overlapped, the determinations were considered equal. \*Positive for PR3-ANCA in at least one antigen-specific immunoassay (*Finkelstein et al. Am J Med.* 2007); \*\*Positive for PR3-ANCA in at least one antigen-specific immunoassay (*Fussner et al. Arthritis Rheum* 2016)].

**Fig. 2.**
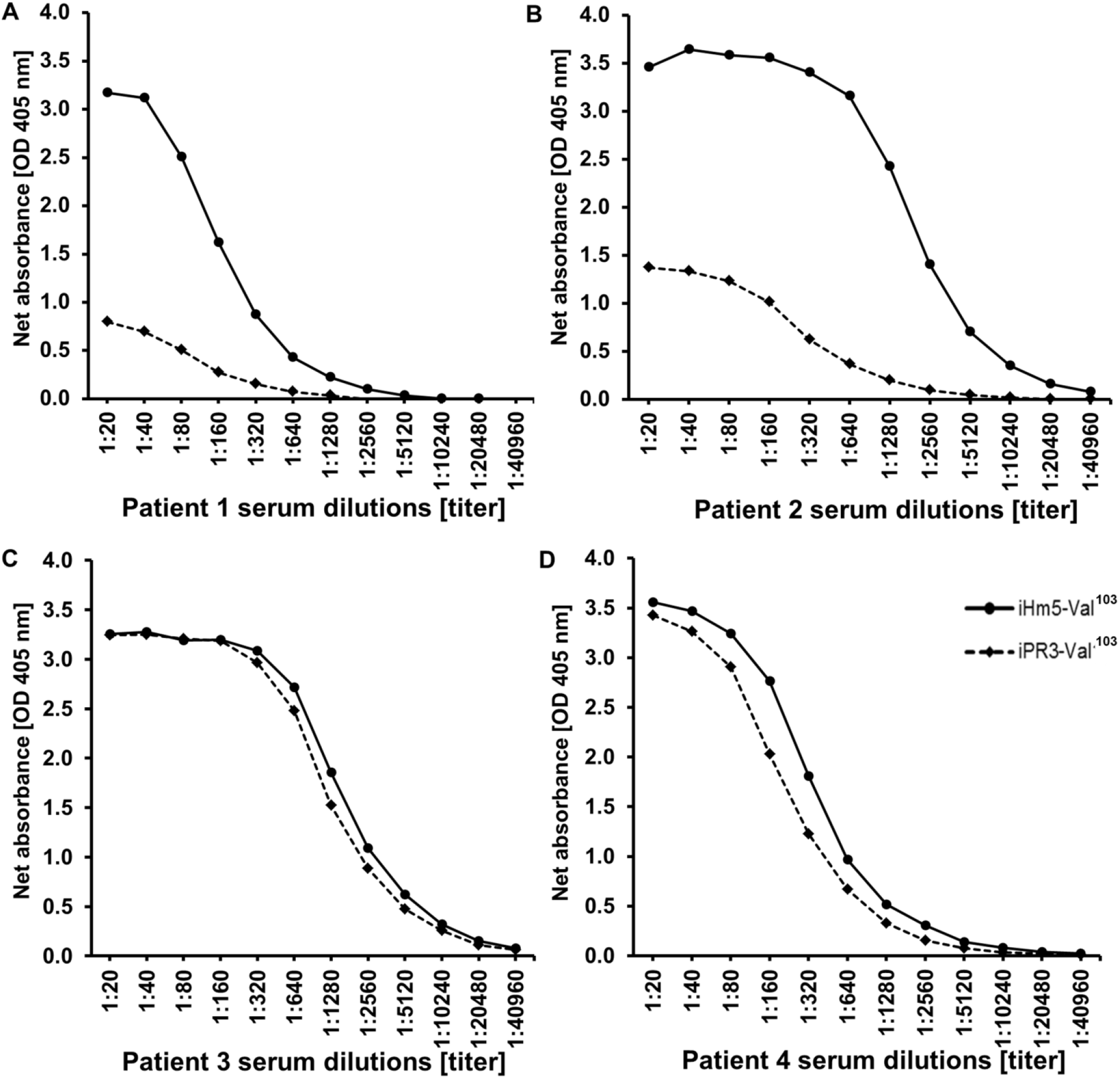
Reactivity of PR3-ANCA positive serum samples with iHm5 and iPR3. Representative examples of serum samples from patients with PR3-ANCA associated vasculitis showing higher (A and B) or equal (C and D) binding to iHm5 compared to iPR3 in the anchor ELISA.

**Fig. 3.**
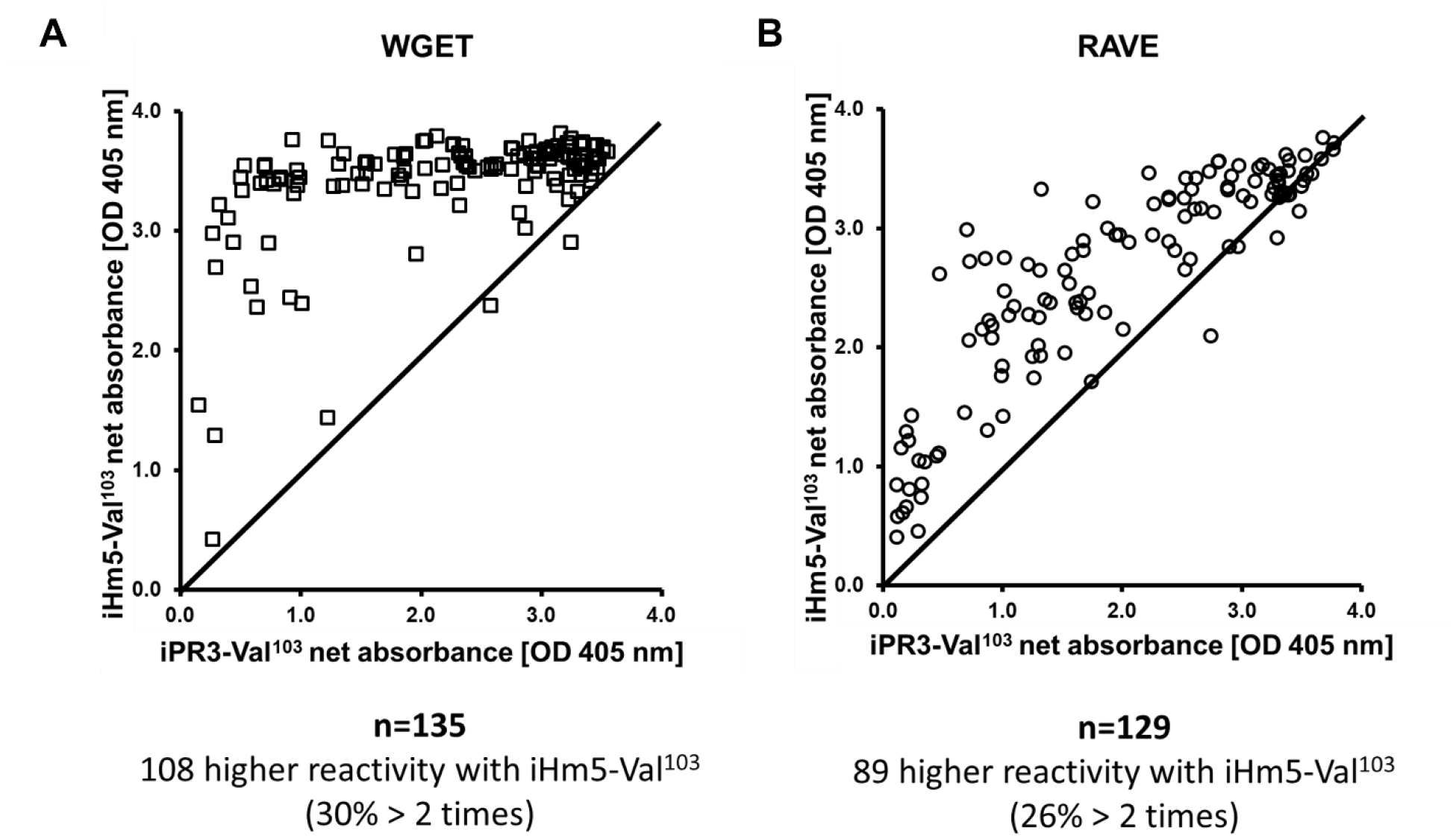
PR3-ANCA reactivity with iHm5 and iPR3 determined by anchor ELISA using serum from patients included in the WGET and in the RAVE trials. Scatter plot of the net absorbance values of iHm5 recognition plotted against iPR3 recognition using the anchor ELISA described with plasma and sera from patients of two AAV cohorts, the WGET (A) and the RAVE (B), respectively. There is a shift of the net absorbance values towards the binding of patient PR3-ANCA to iHm5. (Abbreviations: OD - optical density; PR3 - proteinase 3; RAVE – rituximab vs. cyclophosphamide for remission induction in AAV; WGET - Etanercept vs. placebo for remission induction in AAV)

To confirm these results in an independent cohort, we also analyzed 129 serum samples, which had previously tested positive for PR3-ANCAs in at least one immunoassay and were obtained from the participants in the RAVE study at the time of enrollment (35, 36). Of these, 128 samples showed reactivity with iHm5 and 126 samples reacted with both iHm5 and iPR3. The reactivities with iHm5 of 33 (26%) of these 126 samples were more than double the reactivities with iPR3 (**Fig 3 & Table 1**). The single serum sample from the WGET cohort and the three serum samples from the RAVE cohort that showed higher reactivity with iPR3 than with iHm5 all yielded very high net absorbance values for both antigens, and on repeat testing results were consistently slightly lower with iHm5 with the differences for each sample being just above the intra-assay coefficient of variance of the assay of 5% (data not shown).

To further evaluate whether the preferential binding of PR3-ANCA to iHm5 over iPR3 is of potential clinical utility, we explored the sensitivity and specificity of the anchor ELISA using iHm5 in defined cohorts.

Of the 14 WGET participants who had previously consistently tested negative for PR3-ANCAs in other assays (28), enrollment serum samples of two patients (14%) were positive in the anchor ELISA using iHm5, indicating that in a sizable number of patients with GPA that were previously labeled as “ANCA-negative,” PR3-ANCA can be detected with the iHm5 mutant. By contrast, among 52 MPO-ANCA positive participants of the RAVE study, a patient population not expected to have PR3-ANCAs, two (3.8%) serum samples tested positive (net absorbance of 0.625 and 0.125 OD, respectively) in the anchor ELISA using iHm5 and negative when using iPR3.

We also determined the prevalence of PR3-ANCAs in a population of 300 octogenarians without AAV by the anchor ELISA using iHm5 and iPR3 because the frequency of autoantibodies without a corresponding disease is reportedly increasing with age (45). Only one sample from this population showed weak IgG PR3-ANCA reactivity with iHm5 by the anchor ELISA (net absorbance of 0.173 OD), and two different patients showed reactivity using iPR3 (net absorbances of 0.115 and 0.245 OD, respectively).

To determine whether using iHm5 in the anchor ELISA would allow an earlier detection of PR3-ANCA seroconversion during serial follow-up of patients with AAV, we compared the seroconversion patterns of 33 participants from the WGET and found that PR3-ANCA seroconversion detected by the anchor ELISA using iPR3 could not be detected at an earlier quarterly study visit when using iHm5.

We also compared the utility of iHm5 to iPR3 for IgA and IgM PR3-ANCA detections by the anchor ELISA and found similar preferential binding of IgA and IgM PR3-ANCAs to iHm5 (data not shown).

Taken together, these results indicate preferential reactivity of the majority of PR3-ANCA positive serum samples to iHm5 compared with iPR3 and suggest that iHm5 may enable better detection of low-levels of PR3-ANCAs in patients with AAV than iPR3 without significantly increasing the frequency of false positive results in patients without AAV.

### 3.2. Selective binding of moANCA518 to iHm5

Twenty-five human moAbs derived from plasmablasts of patients with PR3-ANCA positive relapsing AAV were screened for binding to ANCA target antigens by the anchor ELISA using iPR3, iHm5, and two other ANCA-targeting antigens (MPO and HNE).

Interestingly, we found that one such moAb (moANCA518) exhibited selective binding to iHm5 in the anchor ELISA and by Western blot (32), whereas none of these moAbs bound to iPR3, MPO, or HNE. Here we confirmed the selective binding of moANCA518 to iHm5 over iPR3 in a direct ELISA (**Supplementary Fig. 1A**). By contrast, murine anti-PR3 moAbs (PR3G-2, MCPR3-3 and WGM2) exhibited equal binding to both antigens in the direct ELISA (**Supplementary Fig. 1B**).

### 3.3. Remote mutation–induced selective binding of PR3-ANCA to Epitope 3 on iHm5

To identify the epitope recognized by moANCA518 on iHm5 in a capture ELISA, we used epitope-specific moAbs to capture iHm5 and block or modulate the binding of moANCA518 to the epitope that the moAbs recognize (19). As previously described by our group, we found that PR3G-2, a moAb that recognizes Epitope 1 of PR3 (43), did not affect binding of moANCA518 to iHm5; whereas MCPR3-3 or WGM2, both of which recognize Epitope 3 of PR3 (19), respectively decreased or abolished the moANCA518 binding (*p* < 0.01; **Fig. 4A**)(32). When using a PR3-ANCA–containing serum sample obtained from the same patient at the same study visit, from which the plasmablast expressing moANCA518 was generated, we observed similar effects of these moAbs on the reactivity of the PR3-ANCA serum sample with iHm5 (**Fig. 4B**). These findings identify Epitope 3 on iHm5 as the epitope for moANCA518 and for a significant proportion of PR3-ANCA present in the serum obtained at the same time as the moANCA518-generating plasmablasts.

**Fig. 4.**
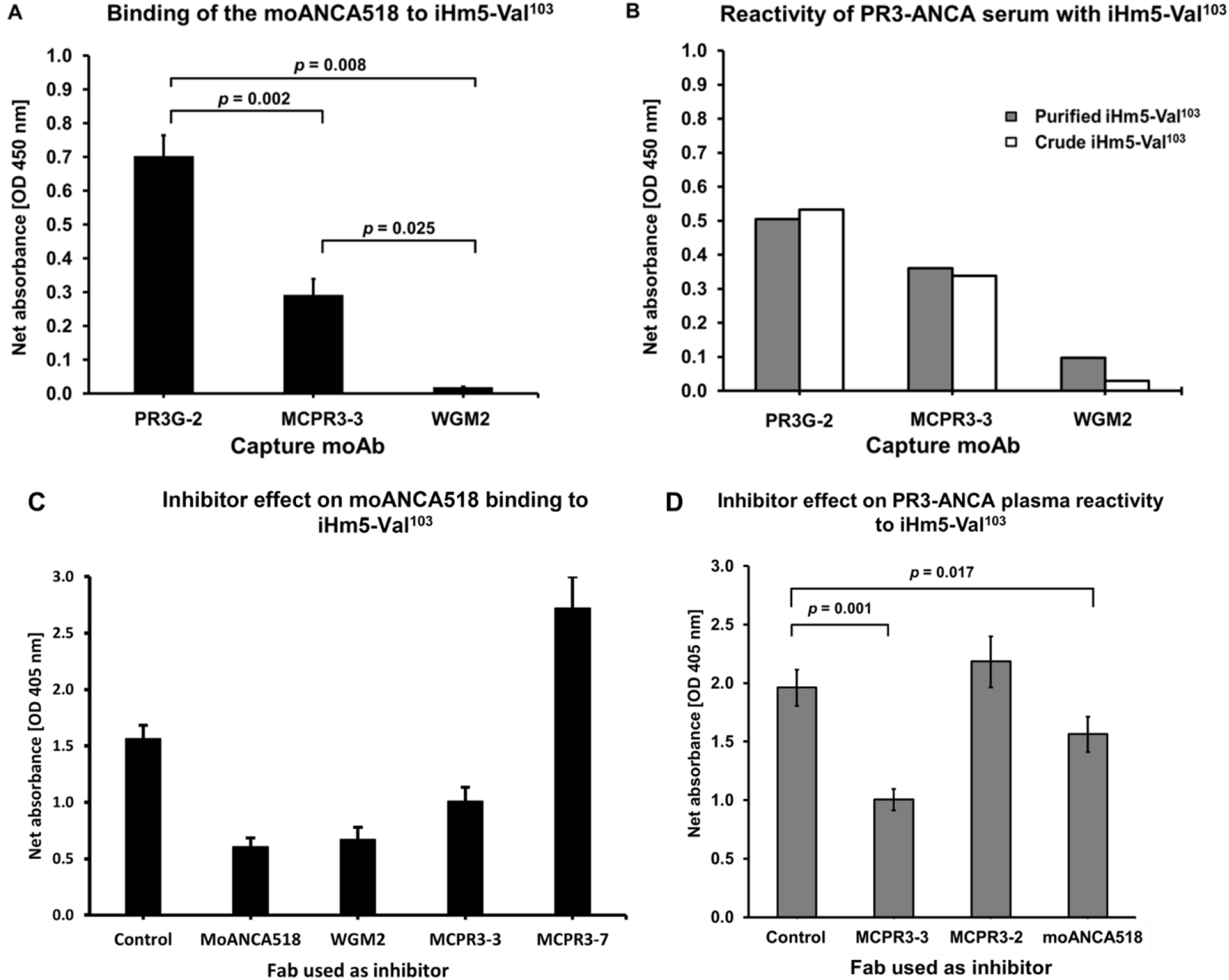
Identification of Epitope 3 of iHm5 as a target of moANCA518 and PR3-ANCA. **A**. Inhibition of the moANCA518 binding to iHm5 using different moAbs was assessed by capture ELISA. The moAbs PR3G-2, MCPR3-3, and WGM2 (2, 4, and 4 µg/mL, respectively) were coated to the plate and used to capture the antigen, iHm5. moANCA518 (1.0 µg/mL) was used as primary antibody. The binding of moANCA518 to iHm5 was impaired significantly when the moAbs MCPR3-3 and WGM2, recognizing Epitope 3, were used to capture the antigen, compared to PR3G-2 which binds to Epitope 1. **B** Serum was obtained from the same patient and study visit from which moANCA518 was developed from isolated plasmablasts. This showed similar differential reactivity of PR3-ANCA with iHm5 depending on the choice of moAb used to capture the antigen. Results were similar when purified iHm5 antigen [black bars] or the crude iHm5 antigen contained in media supernatant from cells expressing the recombinant antigen [white bars] were used. **C** To test our hypothesis using another methodology, we prepared Fabs (2 µg/mL) from epitope specific moAbs as competitors of binding of moANCA518 to iHm5 used as antigen in the anchor ELISA. A significant reduction of binding of moANCA518 to iHm5 was observed after incubation with self (moANCA518 Fab, p=0.005 in comparison with the control) and Fabs that bind to Epitope 3 (MCPR3-3 and WGM2, p=0.05 and p=0.01, respectively, in comparison with the control). There was no effect of incubation with Fabs from MCPR3-7 which binds to Epitope 5 of iPR3. The control Fab corresponds to a mouse IgG1 Fab. (Abbreviations: ELISA – enzyme-linked immunosorbent assay; moAb – monoclonal antibody; TBS – tris-buffered saline) **D** A representative PR3-ANCA positive patient plasma sample was selected from a cohort based on the differential binding results displayed to iHm5 and iPR3 in the anchor ELISA. The control Fab corresponds to a mouse IgG1 Fab. Inhibition of PR3-ANCA binding to iHm5 by Fabs that bind Epitope 3 was observed: MCPR3-3 (p = 0.001, in comparison with the control) and moANCA518 (p = 0.017, in comparison with the control). (Abbreviations: Fab – antigen binding fragment; moAb – monoclonal antibody; OD – optical density)

We further confirmed Epitope 3 of iHm5 as the primary target for moANCA518 using Fabs from epitope-specific moAbs as inhibitors of binding of moANCA518 to iHm5 using the anchor ELISA (**Fig. 4C**). As expected, we observed the strongest inhibition when the Fab from moANCA518 was used to inhibit the moANCA518 binding. A strong inhibition was observed with the Fabs from MCPR3-3 and WGM2 that target Epitope 3 of PR3 (19, 24). In addition, we found no effect of the Fab from MCPR3-7, which recognizes Epitope 5 (19, 31), on the moANCA518 binding.

The above results demonstrate that moANCA518 recognizes Epitope 3 on iHm5. More importantly, because Epitope 3 is on the opposite side of the chimeric mutation sites in Epitopes 1 and 5 on iHm5 (**Fig. 1**) and has the same amino acid sequence as that of Epitope 3 on iPR3, our observation that moANCA518 recognizes Epitope 3 instead of Epitope 1 or 5 further indicates that the moANCA518 binding is conferred by an unexpected increase of the antigenicity in Epitope 3 induced by the distal mutations in Epitopes 1 and 5.

To determine whether other PR3-ANCAs would preferentially bind to Epitope 3 on iHm5 over iPR3 due to the distal mutations in iHm5, we further evaluated serum or plasma samples from patients with AAV that had exhibited significant preferential reactivity with iHm5 over iPR3 as described in Section 3.1, and found that the Fabs from MCPR3-3 and moANCA518 inhibited significantly (*p* = 0.001 and 0.017, respectively) the reactivity of a representative PR3-ANCA-containing plasma sample to iHm5 in the anchor ELISA (**Fig. 4D**).

These results indicate that the preferential reactivity of PR3-ANCAs containing serum or plasma samples with iHm5 can at least in part be explained by some PR3-ANCAs in these samples being sensitive to the change of antigenicity in Epitope 3 of iHm5 induced by the distal mutations, just like moANCA518.

### 3.4. Capture by monoclonal antibody activates latent epitope on iPR3 for PR3-ANCA recognition

Finally, we wanted to know whether binding of PR3 to antibodies can also induce the activation of distant latent epitopes. Therefore, we evaluated the effect of the binding of the moAb MCPR3-2 on the recognition of iPR3 and iHm5 by the moANCA518. MCPR3-2 binds to Epitope 4 and does not compete for epitopes recognized by a sizable proportion of PR3-ANCA from patients with AAV (24). In contrast to the selective binding of the moANCA518, recognition of iHm5 but not iPR3 (Section 3.1) demonstrated in the immunoassays where the PR3 antigen is presented in isolation (anchor ELISA, direct ELISA, Western blot), i.e. not bound to another protein, the capture of both constructs, iHm5 and iPR3 by MCPR3-2 in the capture ELISA induced the recognition of iPR3 by moANCA518 and further enhanced that of iHm5 compared to the anchor ELISA (**Fig. 5A**). The amount of antigen coated available for antibody binding was comparable between constructs in both assays (**Fig. 5B**).

**Fig. 5.**
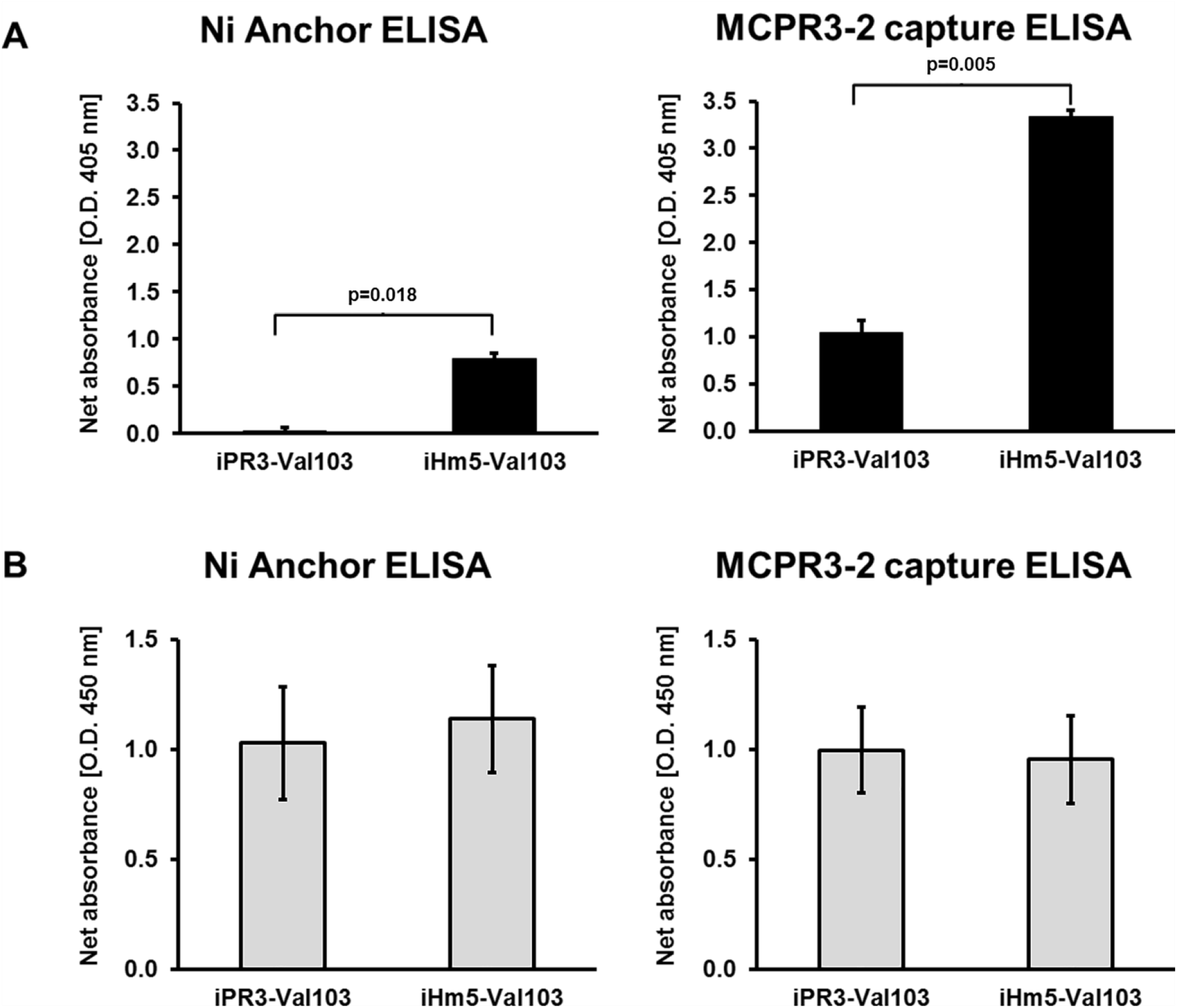
Binding to monoclonal antibodies facilitates iPR3 recognition by the moANCA518. **A** Detection of the binding of moANCA518 to iHm5 and iPR3 using anchor ELISA and MCPR3-2 capture ELISA. Binding of the moANCA518 to iHm5 was detected by anchor ELISA, whereas no binding to iPR3 could be detected (p=0.018). In contrast, on the MCPR3-2 capture ELISA, the moANCA518 bound to both constructs, iHm5 and iPR3, but the signal of the binding to iHm5 was higher when compared with the signal of the binding to iPR3 (p=0.005). **B** Control experiments documenting comparable coating of iPR3 and iHm5, on the anchor ELISA, and comparable coating of MCRP3-2 bound to iPR3 and iHm5, on the MCPR3-2 capture ELISA (p=ns) detected by polyclonal rabbit anti-PR3 antibody. (Abbreviations: ELISA - enzyme-linked immunosorbent assay; PR3 - proteinase 3)

These results indicate that the binding of iPR3 to proteins, in this case to the moAB MCPR3-2 (binding to Epitope 4) can induce mobility changes in Epitope 3 of iPR3 required for the binding of the moANCA518 that mimic the effect of the distant three point mutations of iHm5 on Epitope 3, which favor the recognition by the moANCA518 and by a sizable proportion of PR3-ANCA.

## 4. Discussion

In this study we show that stronger binding of PR3-ANCAs to PR3 can be induced by the activation of a latent epitope caused by a distant mutation or binding of a moAb to a distant epitope. About half of all PR3-ANCA positive sera displayed stronger binding to iHm5 than to iPR3, indicating that iHm5 has its clinical relevance as an antigen for more sensitive immunoassays for PR3-ANCA detection relative to the use of iPR3. Using iHm5 we had discovered a human monoclonal PR3-ANCA, moANCA518, that binds preferentially to Epitope 3 on iHm5 over iPR3 as a consequence of an unexpected increase of the antigenicity in Epitope 3 of iHm5 that was induced by the three remote chimeric mutations in Epitopes 1 and 5. Furthermore, we found that the majority of PR3-ANCA–positive serum or plasma samples from patients with AAV with higher reactivity to iHm5 than to iPR3 also displayed similar preferential binding to Epitope 3 of iHm5 as moANCA518. Finally, the effect of the mutations causing activation of a latent PR3 epitope could be emulated by capturing iPR3 with the Epitope 4 - specific moAB MCPR3-2 as this induced binding of moANCA518 to iPR3.

The PR3 variant iPR3 has been used for two decades as a standard PR3 antigen for sensitive and specific PR3-ANCA detection by immunoassay (7, 22, 24-29). The iHm5 variant containing three point mutations and its “inverse”-chimeric counterpart, iMh5 (formerly referred to as Mh5) were originally designed for discontinuous epitope mapping and structure-function analysis studies (19). When we used iHm5 for the screening of anti-PR3 moAbs and PR3-ANCAs, we expected that some PR3-ANCA positive sera would display reduced binding to iHm5 which would have allowed the conclusion that some PR3-ANCAs in such a serum sample target Epitope 5. Instead, we identified no loss of binding to iHm5 compared to iPR3 by any PR3-ANCA positive sera from patients with PR3-ANCA positive AAV, but preferential reactivity of the majority of PR3-ANCA positive serum samples to iHm5 compared with iPR3 indicating that iHm5 may enable better detection of low-levels of PR3-ANCAs in patients with AAV than iPR3. Consequently, we investigated whether iHm5 represents a useful antigen for more sensitive immunoassays for PR3-ANCA detection. We found that low-level PR3-ANCAs can be detected more readily in some patients with PR3-ANCA positive AAV without generating a significant number of false positive results in patients with MPO-ANCA positive vasculitis or octogenarians without AAV. However, we were unable to detect recurrent PR3-ANCAs earlier using iHm5 versus iPR3 as antigen when testing the serial follow-up serum samples obtained from trial participants at quarterly intervals (34-36). Whether the use of iHm5 might be clinically useful for early detection of PR3-ANCA seroconversion after rituximab therapy in chronically relapsing patients with PR3-ANCA associated AAV, would require a dedicated prospective study with PR3-ANCA measurements at narrower intervals.

The identification of moANCA518 with selective binding to iHm5 was a fortuitous finding that allowed us to further investigate the observation of increased reactivity of many PR3-ANCA–positive sera with iHm5. The PR3-ANCA response is thought to be an oligoclonal immune response as serum samples from patients have been documented to contain PR3-ANCAs binding to more than one epitope (19, 46). The systematic analysis of the binding of moANCA518 to iHm5 using moAbs with defined PR3-ANCA epitope recognition as inhibitors of binding led to another surprise, namely, that moANCA518 does not bind to the surface region of PR3 where the mutations that differentiate iHm5 from iPR3 are located, but to Epitope 3 of iHm5 which is located on the opposite side of the iHm5 structure from the side where the mutations were introduced (**Fig. 1**) (32). This allowed the use of moANCA518 binding to iHm5 and iPR3 as a gauge to determine that serum PR3-ANCA from patients do indeed contain antibodies that can only bind to PR3 when the latent epitope is activated.

It has previously been described for antiphospholipid syndrome that changes in the conformation of the target antigen induced by the binding to cardiolipin vesicles result in the exposure of previously inaccessible epitopes rendering them available for recognition by pathogenic autoantibodies (47). Here, we demonstrated that the binding of the moAb MCPR3-2 to iPR3 on Epitope 4 induced recognition of iPR3 by the moANCA518. This finding has two implications. First, it may explain why the MCPR3-2 capture ELISA method for PR3-ANCA detection has consistently been more sensitive than other methods without loss of specificity as some PR3-ANCA can only bind if a latent epitope is activated by an antibody-antigen interaction. Second, it implies a previously unrecognized level of complexity and variability of the oligoclonal PR3-ANCA interactions with its target antigen in individual patients. Consequently, one can assume that the functional impact including pathogenicity of PR3-ANCA may change as the components of the oligoclonal mixture change in patients over time.

In conclusion, this study shows that iHm5 is a clinically relevant PR3 mutant which can be used as an antigen for sensitive PR3-ANCA detection in patients with AAV without significant false positive PR3-ANCA detection in patients without AAV. More importantly, our results demonstrate that the preferential binding of PR3-ANCAs to iHm5 over iPR3 is the result of an increase of the antigenicity of Epitope 3 on iHm5 induced by the three distal point mutations, and a similar effect can be induced in Epitope 3 by engagement of the separate Epitope 4 by a moAB. Consequently, the present work suggests that remote activation or potentiation of a latent epitope can be achieved not only by distant point mutations but also by the binding of antibodies or possibly other ligands to PR3. Further investigations are needed to determine whether these mechanisms play a role in etiologies of antibody-mediated autoimmune diseases and whether they can be utilized for the development of novel treatment strategies for these diseases.

## Acknowledgements

The moAbs MCPR3-3 and MCPR3-7 were generated with the assistance of Thomas G. Beito from the Mayo Clinic Monoclonal Antibody Core Facility.

### Appendix A – Supplementary data

#### Supplementary Figure

**Supplementary Fig. 1.**
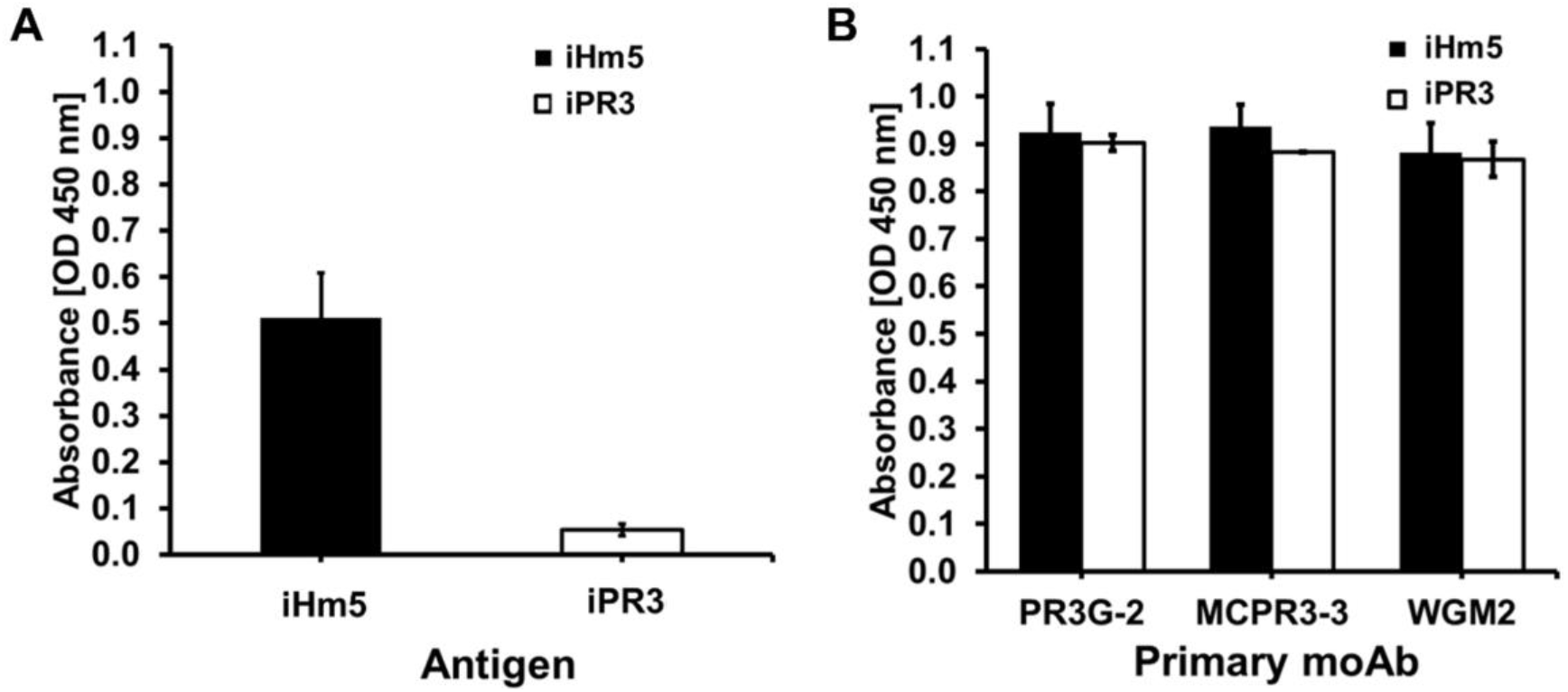
Selective binding of moANCA518 to iHm5 over iPR3. **A** Use of the antigens iHm5 and iPR3 in a direct ELISA showed binding of moANCA518 to iHm5 (black bar) but not to iPR3 (white bar). **B**. Binding of the mouse moAbs PR3G-2, MCPR3-3, and WGM2 to iHm5 (black bars) and iPR3 (white bars) in the direct ELISA. The absorbances at 450 nm for the binding to iHm5 and iPR3 were similar for all moAbs. (Abbreviations: ELISA – enzyme-linked immunosorbent assay; HRP - horseradish peroxidase; moAb – monoclonal antibody.

## References

1. Jennette JC, Falk RJ, Bacon PA, Basu N, Cid MC, Ferrario F, et al. 2012 revised international Chapel Hill consensus conference nomenclature of vasculitides. Arthritis Rheumatol 2013;65:1–11.

2. Falk RJ, Jennette JC. Anti-neutrophil cytoplasmic autoantibodies with specificity for myeloperoxidase in patients with systemic vasculitis and idiopathic necrotizing and crescentic glomerulonephritis. N Engl J Med 1988;318:1651–1657.

3. Jenne DE, Tschopp J, Lüdemann J, Utecht B, Gross WL. Wegener’s autoantigen decoded. Nature 1990;346:520.

4. Hoffman GS, Specks U. Antineutrophil cytoplasmic antibodies, Arthritis Rheumatol 1998;41:1521–1537.

5. Bossuyt X, Cohen Tervaert JW, Arimura Y, Blockmans D, Flores-Suarez LF, Guillevin L, et al. Position paper: revised 2017 international consensus on testing of ANCAs in granulomatosis with polyangiitis and microscopic polyangiitis. Nat Rev Rheumatol 2017;13:683–692.

6. Cornec D, Cornec-Le Gall E, Fervenza FC, Specks U. ANCA-associated vasculitis - clinical utility of using ANCA specificity to classify patients. Nat Rev Rheumatol 2016;12:570–579.

7. Fussner LA, Hummel AM, Schroeder DR, Silva F, Cartin-Ceba R, Snyder MR, et al. Factors determining the clinical utility of serial measurements of antineutrophil cytoplasmic antibodies targeting proteinase 3. Arthritis Rheumatol 2016;68:1700–1710.

8. Kemna MJ, Damoiseaux J, Austen J, Winkens B, Peters J, van Paassen P, et al. ANCA as a predictor of relapse: useful in patients with renal involvement but not in patients with nonrenal disease. J Am Soc Nephrol 2015;26:537–542.

9. Kallenberg CG. Pathogenesis of PR3-ANCA associated vasculitis. J Autoimmun 2008;30:29–36.

10. Schönermarck U, Csernok E, Gross WL. Pathogenesis of anti-neutrophil cytoplasmic antibody-associated vasculitis: challenges and solutions 2014. Nephrol Dial Transplant 2015;30 Suppl 1:i46–52.

11. Hutton HL, Holdsworth SR, Kitching AR. ANCA-associated vasculitis: pathogenesis, models, and preclinical testing. Semin Nephrol 2017;37:418–435.

12. Bini P, Gabay JE, Teitel A, Melchior M, Zhou JL, Elkon KB. Antineutrophil cytoplasmic autoantibodies in Wegener’s granulomatosis recognize conformational epitope(s) on proteinase 3. J Immunol 1992;149:1409–1415.

13. van der Geld YM, Oost-Kort W, Limburg PC, Specks U, Kallenberg CG. Recombinant proteinase 3 produced in different expression systems: recognition by anti-PR3 antibodies. J Immunol Methods 2000;44:117–131.

14. Specks U. What you should know about PR3-ANCA conformational requirements of proteinase 3 (PR3) for enzymatic activity and recognition by PR3-ANCA. Arthritis Res 2000;2:263–267.

15. Williams Jr RC, Malone CC, Payabyab J, Byres L, Underwood D. Epitopes on proteinase-3 recognized by antibodies from patients with Wegener’s granulomatosis. J Immunol 1994;152:4722–4737.

16. van der Geld YM, Limburg PC, Kallenberg CG. Proteinase 3, Wegener’s autoantigen: from gene to antigen. J Leukoc Biol 2001;69:177–190.

17. Griffith ME, Coulthart A, Pemberton S, George AJT, Pusey CD. Anti-neutrophil cytoplasmic antibodies (ANCA) from patients with systemic vasculitis recognize restricted epitopes of proteinase 3 involving the catalytic site. Clin Exp Immunol 2001;123:170–177.

18. Bruner BF, Vista ES, Wynn DM, Harley JB, James JA. Anti-neutrophil cytoplasmic antibodies target sequential functional proteinase 3 epitopes in the sera of patients with Wegener’s granulomatosis, Clin Exp Immunol 2010;162:262–270.

19. Silva F, Hummel AM, Jenne DE, Specks U. Discrimination and variable impact of ANCA binding to different surface epitopes on proteinase 3, the Wegener’s autoantigen. J Autoimmun 2010;35:299–308.

20. Gencik M, Meller S, Borgmann S, Fricke H. Proteinase 3 gene polymorphisms and Wegener’s granulomatosis. Kidney Int 2000;58:2473–2477.

21. Dolman KM, Jager A, Sonnenberg A, von dem Borne A, Goldschmeding R. Proteolysis of classic anti-neutrophil cytoplasmic autoantibodies (C-ANCA) by neutrophil proteinase 3. Clin Exp Immunol 1995;101:8–12.

22. Specks U, Fass DN, Fautsch MP, Hummel AM, Viss MA. Recombinant human proteinase 3, the Wegener’s autoantigen, expressed in HMC-1 cells is enzymatically active and recognized by c-ANCA. FEBS Letters 1996;390:265–270.

23. Sun J, Fass DN, Viss MA, Hummel AM, Tang H, Homburger HA, Specks U. A proportion of proteinase 3 (PR3)-specific anti-neutrophil cytoplasmic antibodies (ANCA) only react with PR3 after cleavage of its N-terminal activation dipeptide. Clin Exp Immunol 1998;114:320–326.

24. Sun J, Fass DN, Hudson JA, Viss MA, Wieslander J, Homburger HA, Specks U. Capture-ELISA based on recombinant PR3 is sensitive for PR3–ANCA testing and allows detection of PR3 and PR3–ANCA rPR3 immunecomplexes. J Immunol Methods 1998;211:111–123.

25. Russell KA, Wiegert E, Schroeder DR, Homburger HA, Specks U. Detection of anti-neutrophil cytoplasmic antibodies under actual clinical testing conditions. Clin Immunol 2002;103:196–203.

26. Capizzi SA, Viss M, Hummel AM, Fass DN, Specks U. Effects of carboxy-terminal modifications of proteinase 3 (PR3) on the recognition by PR3-ANCA. Kidney Int 2003;63:756–760.

27. Lee AS, Finkielman JD, Peikert T, Hummel AM, Viss MA, Specks U. A novel capture-ELISA for detection of anti-neutrophil cytoplasmic antibodies (ANCA) based on c-myc peptide recognition in carboxy-terminally tagged recombinant neutrophil serine proteases. J Immunol Methods 2005;307:62–72.

28. Finkielman JD, Lee AS, Hummel AM, Viss MA, Jacob GL, Homburger HA, et al. ANCA are detectable in nearly all patients with active severe Wegener’s granulomatosis. Am J Med 2007;120:643e9–643e14.

29. Finkielman JD, Merkel PA, Schroeder DR, Hoffman GS, Spiera R, St. Clair EW, et al. for the WGET Research Group. Antiproteinase 3 Antineutrophil Cytoplasmic Antibodies and Disease Activity in Wegener Granulomatosis. Ann Intern Med 2007;147:611–619.

30. Korkmaz B, Kuhl A, Bayat B, Santoso S, Jenne DE. A hydrophobic patch on proteinase 3, the target of autoantibodies in Wegener granulomatosis, mediates membrane binding via NB1 Receptors. J Biol Chem 2008;283:35976–35982.

31. Hinkofer LC, Seidel SA, Korkmaz B, Silva F, Hummel AM, Braun D, et al. A monoclonal antibody (MCPR3-7) interfering with the activity of proteinase 3 by an allosteric mechanism. J Biol Chem 2013;288:26635–26648.

32. Pang YP, Casal Moura M, Thompson GE, Nelson DR, Hummel AM, Jenne DE, et al. Remote Activation of a Latent Epitope in an Autoantigen Decoded With Simulated B-Factors. Front Immunol. 2019;10:2467

33. Oommen E, Hummel AM, Allmannsberger L, Cuthbertson D, Carette S, Pagnoux C, et al. IgA antibodies to myeloperoxidase in patients with eosinophilic granulomatosis with polyangiitis (Churg-Strauss). Clin Exp Rheumatol 2017;35:98–101.

34. The Wegener’s Granulomatosis Etanercept Trial (WGET) research group. Etanercept plus standard therapy for Wegener’s granulomatosis. N Engl J Med 2005;352:351–361.

35. Stone JH, Merkel PA, Spiera R, Seo P, Langford CA, Hoffman GS, et al. Rituximab versus cyclophosphamide for ANCA-associated vasculitis. N Engl J Med 2010;363:221–232.

36. Specks U, Merkel PA, Seo P, Spiera R, Langford CA, Hoffman GS, et al. Efficacy of remission-induction regimens for ANCA-associated vasculitis. N Engl J Med 2013;369:417–427.

37. Lee AS, Finkielman JD, Peikert T, Hummel AM, Viss MA, Jacob GL, et al. Agreement of anti-neutrophil cytoplasmic antibody measurements obtained from serum and plasma. Clinical and experimental immunology. 2006;146(1):15–20.

38. Tan YC, Kongpachith S, Blum LK, Ju CH, Lahey LJ, Lu DR, et al. Barcode-enabled sequencing of plasmablast antibody repertoires in rheumatoid arthritis. Arthritis Rheumatol 2014;66:2706–2715.

39. Hershberg U, Meng W, Zhang B, Haff N, St Clair EW, Cohen PL, et al. Persistence and selection of an expanded B-cell clone in the setting of rituximab therapy for Sjögren’s syndrome. Arthritis Res Ther 2014;16:R51.

40. Peikert T, Finkielman JD, Hummel AM, McKenney ME, Gregorini G, Trimarchi M, et al. Functional characterization of antineutrophil cytoplasmic antibodies in patients with cocaine-induced midline destructive lesions. Arthritis Rheumatol 2008;58:1546–1551.

41. Wiesner O, Russell KA, Lee AS, Jenne DE, Trimarchi M, Gregorini G, et al. Antineutrophil cytoplasmic antibodies reacting with human neutrophil elastase as a diagnostic marker for cocaine-induced midline destructive lesions but not autoimmune vasculitis, Arthritis Rheumatol 2004;50:2954–2965.

42. Wiesner O, Litwiller RD, Hummel AM, Viss MA, McDonald CJ, Jenne DE, et al. Differences between human proteinase 3 and neutrophil elastase and their murine homologues are relevant for murine model experiments. FEBS Lett 2005;579:5305–5312.

43. van der Geld YM, Limburg PC, Kallenberg CG. Characterization of monoclonal antibodies to proteinase 3 (PR3) as candidate tools for epitope mapping of human anti-PR3 autoantibodies. Clin Exp Immunol 1999;118:487–496.

44. Csernok E, Lüdemann J, Gross WL, Bainton DF. Ultrastructural localization of proteinase 3, the target antigen of anti-cytoplasmic antibodies circulating in Wegener’s granulomatosis. Am J Pathol 1990;137:1113–1120.

45. Andersen-Ranberg K, HØier-Madsen M, Wiik A, Jeune B, Hegedus L. High prevalence of autoantibodies among Danish centenarians. Clin Exp Immunol 2004;138:158–163.

46. van der Geld YM, Stegeman CA, Kallenberg CG. B cell epitope specificity in ANCA-associated vasculitis: does it matter?. Clin Exp Immunol 2004;137:451–459.

47. de Laat B, Derksen RH, van Lummel M, Pennings MT, deGroot PG. Blood 2006;107(5):1916–1924.

